# Activating mGlu_3_ metabotropic glutamate receptors rescues schizophrenia-like cognitive deficits through metaplastic adaptations within the hippocampus

**DOI:** 10.1101/2020.10.27.356196

**Authors:** Shalini Dogra, Branden J. Stansley, Zixiu Xiang, Weilun Qian, Rocco G. Gogliotti, Ferdinando Nicoletti, Craig W. Lindsley, Colleen M. Niswender, Max E. Joffe, P. Jeffrey Conn

## Abstract

**Background:** Polymorphisms in *GRM3*, the gene encoding the mGlu3 metabotropic glutamate receptor, are associated with impaired cognition and neuropsychiatric disorders such as schizophrenia. Limited availability of selective genetic and molecular tools has hindered progress in developing a clear understanding of the mechanisms through which mGlu3 receptors regulate synaptic plasticity and cognition.

**Methods:** We examined associative learning in mice with trace fear conditioning, a hippocampal-dependent learning task disrupted in patients with schizophrenia. Underlying cellular mechanisms were assessed using *ex vivo* hippocampal slice preparations with selective pharmacological tools and selective genetic deletion of mGlu3 receptor expression in specific neuronal subpopulations.

**Results:** mGlu_3_ receptor activation enhanced trace fear conditioning and reversed deficits induced by subchronic phencyclidine. Mechanistic studies revealed that mGlu_3_ receptor activation induced metaplastic changes, biasing afferent stimulation to induce long-term potentiation through a mGlu_5_ receptor-dependent, endocannabinoid-mediated, disinhibitory mechanism. Selective genetic deletion of either mGlu_3_ or mGlu_5_ from hippocampal pyramidal cells eliminated effects of mGlu_3_ activation, revealing a novel mechanism by which mGlu_3_ and mGlu_5_ interact to enhance cognitive function.

**Conclusions:** These data demonstrate that activation of mGlu_3_ receptors in hippocampal pyramidal cells enhances hippocampal-dependent cognition in control and impaired mice by inducing a novel form of metaplasticity to regulate circuit function – providing a clear mechanism through which genetic variation in *GRM3* can contribute to cognitive deficits. Developing approaches to positively modulate mGlu_3_ receptor function represents an encouraging new avenue for treating cognitive disruption in schizophrenia and other psychiatric diseases.

## Introduction

Recent studies have identified disease-related mutations associated with cognitive deficits in patients with schizophrenia (1). Several analyses, including a 37,000-patient genome-wide association study, have implicated single nucleotide polymorphisms in the *GRM3* gene with an increased likelihood of schizophrenia diagnosis (2–4). Moreover, studies have linked *GRM3* variation with performance on memory-based tasks in schizophrenia patients and neurotypical controls (5–7). *GRM3* codes for the mGlu_3_ subtype of metabotropic glutamate receptor. One well-studied *GRM3* polymorphism is associated with splice variants that translate to truncated, non-functional mGlu_3_ receptor protein (8, 9), suggesting decreased mGlu_3_ receptor function contributes to schizophrenia etiology in some individuals. Preclinical studies bolster the clinical association between mGlu_3_ receptor function and cognition. In mice, for example, global deletion of *Grm3* induces deficits in reference memory and working memory (10, 11), but the circuits and molecules that mediate this phenotype remain unresolved.

While mouse and human genetic studies suggest a relationship between *GRM3* and schizophrenia, the paucity of selective tools has hindered the clear delineation of mechanisms that link mGlu_3_ receptor function and hippocampal cognition. A large body of preclinical work has prompted great interest in targeting mGlu_3_ (and mGlu_2_) receptors for the treatment of schizophrenia (reviewed in (3)). This literature is exemplified by seminal work from Moghaddam and Adams (12), which first indicated that non-selective orthosteric mGlu_2/3_ agonists reverse psychotomimetic effects of acute phencyclidine (PCP) administration. Efforts in developing efficacious mGlu_2/3_ agonists culminated in a mixture of promising and disappointing phase II clinical trials. These trials were generally motivated by research showing that mGlu_2_ receptor activation attenuates behaviors that model positive symptoms and psychosis (13–16). On the other hand, mGlu_2_ receptor agonism suppresses rapid eye movement sleep (15) and can impair learning and memory on hippocampal-dependent tasks (17). These findings suggest that, in clinical trials with non-selective compounds, activating mGlu_2_ receptors may have induced cognitive side effects that obscured latent benefits of mGlu_3_ receptor activation. Collectively, the clinical and translational findings with mGlu_2/3_ agonists, along with genetic studies implicating *GRM3*, raise the exciting possibility that selective activation of mGlu_3_ could improve cognition, particularly in models of schizophrenia-like deficits.

Here, we leverage the recent development of highly selective allosteric modulators to demonstrate that mGlu_3_ receptor activation enhances cognition. Moreover, we report that mGlu_3_ receptors regulate hippocampal plasticity by a novel mechanism that requires co-activation of a second mGlu receptor, mGlu_5_. Finally, using newly developed genetic mouse models to selectively delete mGlu_3_ and mGlu_5_ from pyramidal cells, we relate these changes in synaptic plasticity to changes in associative learning. These data build on exciting advances in human genetics and reveal mGlu_3_ receptors as novel targets for ameliorating cognitive symptoms in schizophrenia.

## Methods and Materials

### Animals

Mice were cared for in accordance with the National Institutes of Health *Guide for the Care and Use of Laboratory Animals*. Studies were approved by the Vanderbilt University School of Medicine IACUC and occurred during the light phase. Young adult (6-8-week-old) C57BL/6J (Jackson Laboratories), *Grm5*^Fl/Fl^, *Grm5*-CaMKII KO, *Grm3*^*Fl/Fl*^, *Grm3*-CMV KO, *and Grm3-*CaMKII KO mice (Key Resources Table) were housed on a standard light cycle (on at 6:00) and provided *ad libitum* food and water. Strains were backcrossed to congenic C57BL/6J mice for more than 5 generations. Both male and female mice were used.

### Drugs

Most drugs were purchased from Tocris (Key Resources Table). PCP was purchased from Sigma. VU0469650, VU0650786, VU6001966 were synthesized in-house (18–20). Test compounds were delivered by intraperitoneal injection (10% Tween-80, 10 μL/kg).

### Subchronic phencyclidine (scPCP) treatment

5-week-old mice received daily subcutaneous injections with 10 mg/kg PCP (0.9% saline, 10 μL/kg) for 7 days. Separate cohorts of mice were used for electrophysiology or behavior following a 7-day washout period.

### Extracellular field potential recordings

Experiments were performed as described previously (21, 22). In brief, mice were anesthetized with isoflurane and coronal slices (400 μm) containing the dorsal hippocampus were prepared in *N*-methyl-D-glucamine-based solution. Slices were held at room temperature in ACSF containing (in mM): 126 NaCl, 1.25 NaH_2_PO_4_, 2.5 KCl, 10 D-glucose, 26 NaHCO_3_, 2 CaCl_2_, and 1 MgSO_4_. We recorded field excitatory postsynaptic potentials (fEPSPs) with glass electrodes (1-3 MΩ) in the stratum radiatum of CA1 and a bipolar stimulating electrode near the CA3-CA1 border. Stimuli were constantly delivered at 0.05 Hz unless otherwise noted. Theta burst stimulation (TBS) long-term potentiation (LTP) was evoked using 9 bursts of 4 stimuli at 100 Hz, repeated every 230 ms (21). Afferent ‘priming’ consisted of 2 bursts of 10 stimuli at 10 Hz, separated by 20 seconds (23). Long-term depression (LTD) was evoked with DHPG or with 15 minutes of paired pulse 1 Hz stimulation, each in the presence of the mGlu_1_ NAM VU0469650 (10 μM). Data were digitized using a Multiclamp 700B, Digidata 1322A, and pClamp 10 software (Molecular Devices). Slices from at least 3 mice are included in each group.

### Whole-cell voltage-clamp recordings

Whole-cell recordings were made from hippocampal CA1 pyramidal neuron somata. The pipette was filled with intracellular solution (mM): 125 CsCl, 4 NaCl, 0.5 MgCl_2_, 10 TEA, 10 HEPES, 0.5 EGTA, 5 QX-314, 5 Tris-phosphocreatine, 4 ATP-Mg and 0.3 GTP-Na, adjusted to pH 7.3 and 290-295 mOsm. Inhibitory postsynaptic currents (IPSCs) were evoked at 0.1 Hz and recorded at −70 mV in CNQX (20 μM) and AP-5 (50 μM).

### Trace fear conditioning

Design for experiments was modified from previous studies (23). Vehicle or LY379268 was injected 30 minutes before the session. All mGlu negative allosteric modulators were injected 20 minutes prior to LY379268/Vehicle treatment. Mice were placed in a conditioning chamber with a shock grid (Med Associates) in the presence of 10% vanilla odor. Mice were acclimated for 60 seconds before conditioning and a 15-second tone (90dB, 2900Hz) was applied preceding a 1-second foot shock (0.5-mA mild shock or 0.7-mA strong shock). A precise 30-second interval, or ‘trace’, separated the tone and shock. Three tone-trace-shock pairings (four pairings for scPCP study) were applied, 240-seconds apart. Freezing was quantified during each trace by video software (VideoFreeze) and confirmed by a blinded observer.

### Generation of floxed *Grm3* mice

We generated *Grm3^Fl/Fl^* mice using embryonic stem cell-mediated gene targeting on the C57BL/6J genetic background (Ingenious Targeting Laboratory) in which exon 3 of *Grm3* is flanked by LoxP sites (Supplemental Methods).

### RNA and cDNA preparation

Tissues were homogenized and total RNA was prepared using standard Trizol-chloroform methodology. RNA concentration was measured using Nanodrop and cDNA was synthesized from 2 μg total RNA with the SuperScript VILO kit (Thermo Fisher). Primers targeted *Grm3*: F 5’-TGTGGTGAATGCAGTGTACG-3’, R 5’-CATCCCGTCTCCAAAAGTGT-3’ and Actin: F 5’-GTGGGCCGCTCTAGGCACCAA-3’, R 5’-CTCTTTGATGTCACGCACGATTTC-3’.

### Western blotting

Tissues were snap-frozen in liquid nitrogen and stored at −80°C. Tissues were homogenized in radioimmunoprecipitation assay buffer containing protease and phosphatase inhibitors (Sigma Aldrich). Homogenized samples were spun at 12000 g at 4°C for 20 minutes and western bots were run using 50 μg of protein electro-transferred to polyvinyl difluoride membrane. mGlu_3_ receptor primary antibody (Key Resources Table) was added overnight at 4°C followed by incubation with fluorescent secondary antibody for 30 minutes at room temperature. mGlu_3_ receptor protein expression was quantified relative to GAPDH and normalized to values from control mice.

### RNAscope *in situ* hybridization

RNAscope was conducted according to the Advanced Cell Diagnostics user manual (Supplemental Methods). Thin brain slices (16 μm) were incubated with cell type-specific probes and with probes we designed to recognize sequences targeted for excision in *Grm3^Fl/Fl^* and *Grm5^Fl/Fl^* mice (Key Resources Table).

### Statistics

Sample sizes were determined based on previous experiments (22). Analyses were performed using GraphPad Prism 8. Data were represent as mean ± SEM. Significance between groups was determined using Student’s t-test or ANOVA with appropriate post-hoc tests, as specified in the figure legends.

## Results

### mGlu_3_ receptor activation enhances trace fear conditioning

Genetic studies have revealed an association between *GRM3* variation and cognition (5–7). We therefore asked whether activating the mGlu_3_ receptor would improve trace conditioning, a high-demand associative learning task disrupted in patients with schizophrenia (24). Mice were placed in fear conditioning chambers and received three pairings of a tone cue, 30-second “inactive” trace, and mild foot shock (Figure 1A). Mice rapidly associate the trace period with the upcoming shock and express this association by conditioned freezing. We activated mGlu_3_ by systemic delivery of the mGlu_2/3_ agonist LY379268 (3 mg/kg). LY379268 treatment enhanced freezing during the traces proceeding shock presentation (Figure 1B). We next separated the contributions of mGlu_2_ and mGlu_3_ receptors using selective negative allosteric modulators (NAMs). NAM doses were selected based on previously published pharmacokinetic (19, 20) and behavioral studies (25, 26). Either the mGlu_2_ NAM VU6001966 (10 mg/kg, *i.p.*) or the mGlu_3_ NAM VU0650786 (30 mg/kg, *i.p.*) was injected 20 minutes prior to LY379268 treatment. Administration of VU6001966 did not block the LY379268-induced increase in trace fear conditioning (Figure 1B). In contrast, administration of the selective mGlu_3_ NAM VU0650786 (30 mg/kg) blocked the LY379268-induced trace fear conditioning enhancement (Figure 1B). No drug combinations affected freezing during the baseline or first trace, indicating specificity for the learned components of the task. Together, these data suggest that mGlu_3_ receptor activation enhances associative learning in mice.

**Figure 1.**
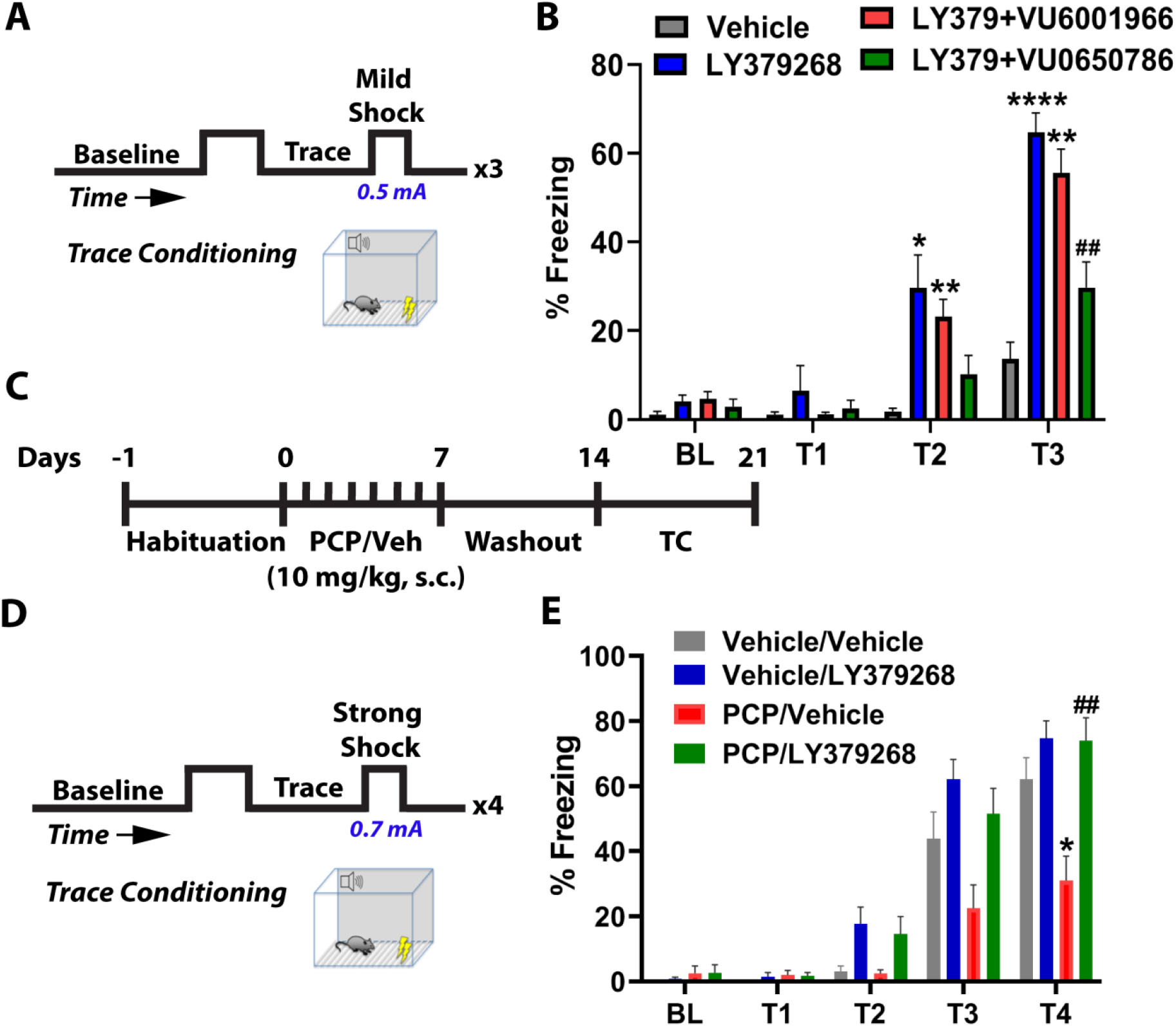
Activation of mGlu_3_ receptors enhances associative learning and rescues schizophrenia-like cognitive deficits. (A) Behavioral schematic. Mice received 3 pairings of a tone and mild foot-shock (0.5 mA), each separated by a 30-second trace period. Freezing time was quantified across the 3 traces. (B) Administration of the mGlu_2/3_ agonist LY379268 (3 mg/kg, *i.p.,* blue bars) 30 minutes prior to conditioning increased freezing at trace 2 and trace 3 relative to vehicle treated mice (grey bars) (*p<0.05, ****p<0.0001). Mice were treated with the mGlu_2_ negative allosteric modulator (NAM) VU6001966 (10 mg/kg, *i.p.*, red bars) or the mGlu_3_ NAM VU0650786 (30 mg/kg, *i.p.*, green bars) to isolate the effects of mGlu_2_ and mGlu_3_ receptor activation. LY379268 enhanced trace conditioning in VU6001966-treated mice (red bars, **p<0.01, compared to vehicle (gray bars)), whereas the effect was blocked by the mGlu_3_ NAM VU0650786 (##p<0.01 compared to LY379268), F(3,72)=37.26, p<0.01, two-way repeated measures ANOVA with Tukey’s post-hoc test, N=11-27 mice). (C) Schematic representing subchronic phencyclidine (scPCP) treatment regimen. After 7 days of habituation, mice were injected with scPCP (10 mg/kg) for 7 days. Trace fear conditioning experiments were performed 7 days after last PCP injection. (D) Behavioral schematic. Mice received 4 pairings of a tone, trace, and strong foot-shock (0.7 mA). (E) scPCP-treated mice (orange bars) froze less than vehicle controls (*p<0.05 compared to vehicle/vehicle; gray bars). Acute LY379268 treatment (30 minutes prior to the experiment; green bars) rescued scPCP-induced deficits in freezing (##p<0.01 compared to PCP/vehicle, red bars), F(3,33)=7.882, p<0.05, two-way repeated measures ANOVA with Tukey’s post-hoc test, N=8-11). Data are presented as mean ± SEM.

### mGlu_3_ agonism rescues PCP-induced deficits in cognition and synaptic plasticity

Subchronic NMDA receptor hypofunction can generate lasting impairments in synaptic plasticity and cognitive function and is thought to model schizophrenia-like forebrain pathophysiology (27, 28). Mice received daily injections of the psychotomimetic NMDA receptor antagonist PCP or Vehicle for one week (scPCP, Figure 1C). Importantly, all treatment groups were tested following a one-week washout, so the behavioral effects of drug combinations do not stem from acute pharmacokinetic or pharmacodynamic interactions with PCP. For this experiment, the number of tone-shock pairings was increased to four and stronger shocks (0.7 mA) were applied to increase freezing in control mice (Figure 1D). Under these conditions, LY379268 no longer enhanced freezing in control mice, likely due to a ceiling effect. Mice treated with scPCP, however, displayed decreased freezing during trace presentations as compared to Vehicle controls. Acute treatment with LY379268 rescued this deficit (Figure 1E), suggesting that mGlu_3_ receptor activation exerts pro-cognitive effects in a model related to schizophrenia-like pathophysiology.

Trace conditioning has been associated with hippocampal function and molecular processes related to synaptic plasticity (28, 29). We therefore evaluated whether scPCP treatment alters LTP at the CA3-CA1 synapse (Figure 2A). TBS induced modest LTP in Vehicle-treated control conditions, but slices from scPCP mice did not display LTP (Figure 2B/2C). Acute slice perfusion with LY379268 (100 nM) completely reversed the plasticity deficit. Together, these findings indicate that mGlu_3_ receptor activation ameliorates coincidental deficits in cognition and hippocampal synaptic plasticity, suggesting that shared molecular mechanisms contribute to both processes. The historical lack of mGlu_3_-selective molecular probes has made it difficult to directly assess effects on synaptic plasticity, thus we performed a series of studies to isolate mGlu_3_ receptors and investigate hippocampal LTP. Acute bath application of the mGlu_2/3_ agonist LY379268 enhanced TBS-LTP in slices from control mice (Figure 2D/2G). The selective mGlu_2_ NAM VU6001966 (10 μM) had no effect on the LY379268-induced increase in LTP (Figure S1). By contrast, the selective mGlu_3_ NAM VU0650786 (20 μM) completely blocked the actions of LY379268 (Figure 2E/2G), confirming that mGlu_3_ receptors are essential for the LTP enhancement.

**Figure 2.**
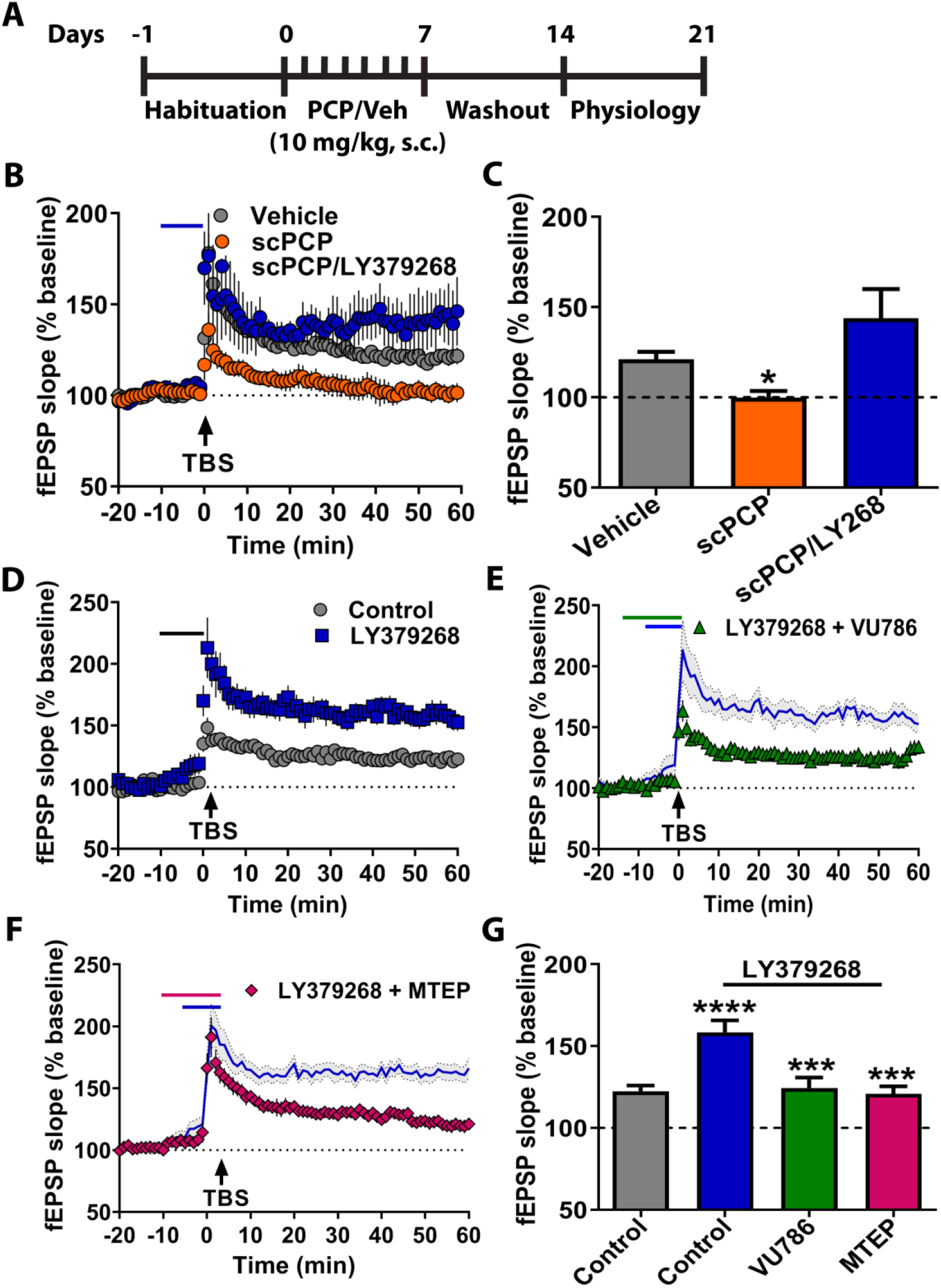
mGlu_3_ receptor activation enhances hippocampal long-term potentiation (LTP) through concerted signaling with mGlu_5_ receptors. (A) Schematic representing scPCP treatment regimen. After 7 days of habituation, mice were injected with scPCP for 7 days. Electrophysiology experiments were performed 7 days after last PCP injection. (B) Field excitatory postsynaptic potentials (fEPSPs) were recorded in the stratum radiatum of CA1 after electrical stimulation of the Schaffer collateral. Time course showing moderate LTP after a single application of theta burst stimulation (TBS) in slices from vehicle-treated mice (grey bar, n=7 slices). Slices from mice treated with scPCP displayed impaired LTP (orange bar n=6). Bath application of the mGlu_2/3_ agonist LY379268 (100 nM) rescued TBS-LTP in slices from scPCP-treated mice (blue bar n=6). (C) Summary of averaged fEPSP slope of last 5 minutes of recordings from panel B (*p<0.05, compared to Vehicle, F(2,16)=5.11, one-way ANOVA with Tukey’s post-hoc test). (D) In control mice, LY379268 application enhanced LTP in response to TBS (blue squares, n=10). (E) Co-application of mGlu_3_ NAM VU0650786 (10 μM) blocked the enhanced LTP induced by LY379268 (green triangles, n=11) and had no effect on its own at baseline. Blue line displays LY379268 data from panel D. (F) The mGlu_5_ NAM MTEP (1 μM) blocked enhanced LTP when co-applied with LY379268 (magenta diamonds, n=8) and had no effect on its own at baseline. (G) Summary of averaged fEPSP slope of last 5 minutes of recordings from panels D-F (****p<0.0001 compared to control, ***p<0.001 compared to LY379268+control, F(3,42)=10.11, one-way ANOVA with Tukey’s post-hoc test). Data are presented as mean ± SEM.

### mGlu_3_ activation enhances LTP through mGlu_5_ receptors

The finding that the mGlu_2/3_ agonist can reverse scPCP-induced deficits in LTP was somewhat surprising since previous studies more commonly implicate group II mGlu receptors in reducing excitatory synaptic transmission. However, previous studies reveal that mGlu_3_ receptors can potentiate or recruit mGlu_5_ receptors (30, 31), which are critical for trace fear conditioning and associative learning (22, 23, 32). We therefore hypothesized that mGlu_3_ agonism potentiates hippocampal LTP through a mechanism that involves co-activation of mGlu_5_ receptors. Consistent with this, the selective mGlu_5_ NAM, MTEP (1 μM), inhibited the ability of LY379268 to potentiate TBS-LTP, without any effect on TBS-LTP alone (Figure 2F/2G). Importantly, all compounds were applied at concentrations that maintain functional selectivity over other mGlu receptor subtypes (31, 33-35). Together, these results suggest that mGlu_3_ receptor activation potentiates TBS-LTP through a mechanism that requires co-activation of mGlu_5_ receptors.

### mGlu_3_ receptor activation biases plasticity towards LTP and away from LTD

We aimed to leverage the extensive literature describing how mGlu_5_ receptors regulate hippocampal plasticity to better understand mGlu_3_ receptor function. The CA3-CA1 synapse is bidirectionally modulated by mGlu_5_ receptors. Modest activation of mGlu_5_ receptors facilitates LTP induction, while stronger activation can promote LTD through distinct signaling cascades (36–38). We reasoned that, if mGlu_3_ receptor activation potentiates all mGlu_5_ receptor-dependent signaling pathways, mGlu_3_ agonists should also enhance LTD. We elicited LTD using the mGlu_1/5_ agonist DHPG (50 μM) co-applied with the mGlu_1_ NAM VU0469650 (10 μM) (Figure 3A/3C). LY379268 (100 nM) impaired LTD (Figure 3B/3C) and did not alter the response to a threshold concentration of DHPG (25 μM) (Figure S2). These data suggest that mGlu_3_ receptor activation induces a qualitative shift in mGlu_5_ receptor signaling to impair LTD. We corroborated these findings using paired-pulse 1-Hz stimulation, which also induces mGlu_5_ receptor-dependent LTD. Remarkably, LY379268 application converted the response to paired-pulse 1-Hz stimulation of Schaffer collateral afferents from induction of LTD to induction of LTP (Figure 3D/3F), fundamentally altering the form of synaptic plasticity induced by low-frequency synaptic activation. This effect was normalized by the mGlu_3_ NAM, VU0650786 (Figure 3E/3F), confirming the involvement of mGlu_3_ receptors. Overall, these results demonstrate that mGlu_3_ activation induces metaplasticity to favor induction of LTP relative to LTD at the CA3-CA1 synapse. We thus aimed to investigate how mGlu_3_ receptors regulate other forms of LTP.

**Figure 3.**
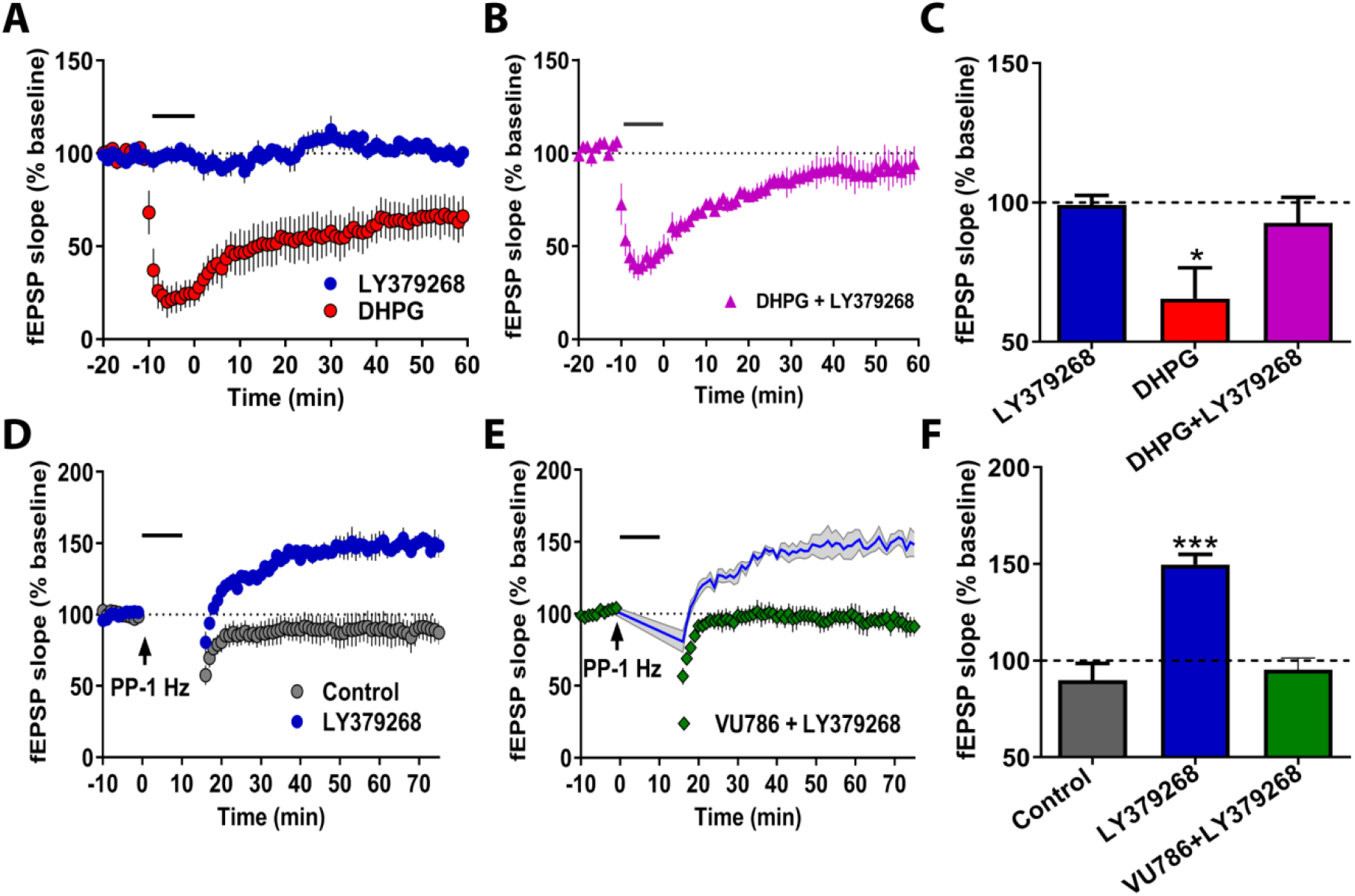
mGlu_3_ receptor activation induces hippocampal metaplasticity to promote LTP. (A) Application of the mGlu_1/5_ agonist DHPG (50 μM) induced long-term depression (LTD) of fEPSP slope (red circles, n=6 slices), while the mGlu_2/3_ agonist LY379268 (100 nM) had no acute or long-term effects (blue circles, n=5). (B) Co-application of LY379268 abrogated DHPG-induced LTD (pink triangles, n=5). (C) Summary of averaged fEPSP slope of last 5 minutes of recordings from panels A-B (*p<0.05, compared to LY379268 alone, F(2,13)=4.087, p<0.05 one-way ANOVA with Tukey’s post-hoc test). (D) Paired-pulse (PP) 1-Hz stimulation for 15 minutes induces mGlu_5_-dependent LTD in control slices (grey circles, n=6). Coincidental application of LY379268 (100 nM) changes PP 1-Hz LTD into a long-term enhancement of fEPSP slope (blue circles, n=6). (E) The mGlu_3_ NAM VU0650786 (10 μM) blocked the metaplastic effects of LY379268 application (green diamonds, n=6). (F) Summary of averaged fEPSP slope of last 5 minutes of recordings from panels D and E (***p<0.001, compared to Control, F(2,13)=17.13, p<0.001, one-way ANOVA with Tukey’s post-hoc test). All experiments were performed in presence of mGlu_1_ NAM, VU0469650 (10 μM). Data are expressed as mean ± SEM.

### mGlu_3_ receptors regulate multiple facets of hippocampal LTP

mGlu receptor activation is necessary and sufficient to induce some forms of LTP at the CA3-CA1 synapse (36, 39–41). Consistent with prior findings, a high concentration of LY379268 (300 nM) induced a slow onset LTP (Figure 4A/4D). Paired-pulse ratios were not altered at any time point (data not shown), suggesting these effects are mediated by postsynaptic mechanisms reminiscent of TBS-LTP. Co-application of either the selective mGlu_3_ NAM VU0650786 (Figure 4B/4D) or the mGlu_5_ NAM, MTEP (Figure 4C/4D) blocked chemically-induced LTP. These data indicate that sustained activation of mGlu_3_ receptors induces LTP, fundamentally contrasting with mGlu_5_ receptor-dependent LTD.

**Figure 4.**
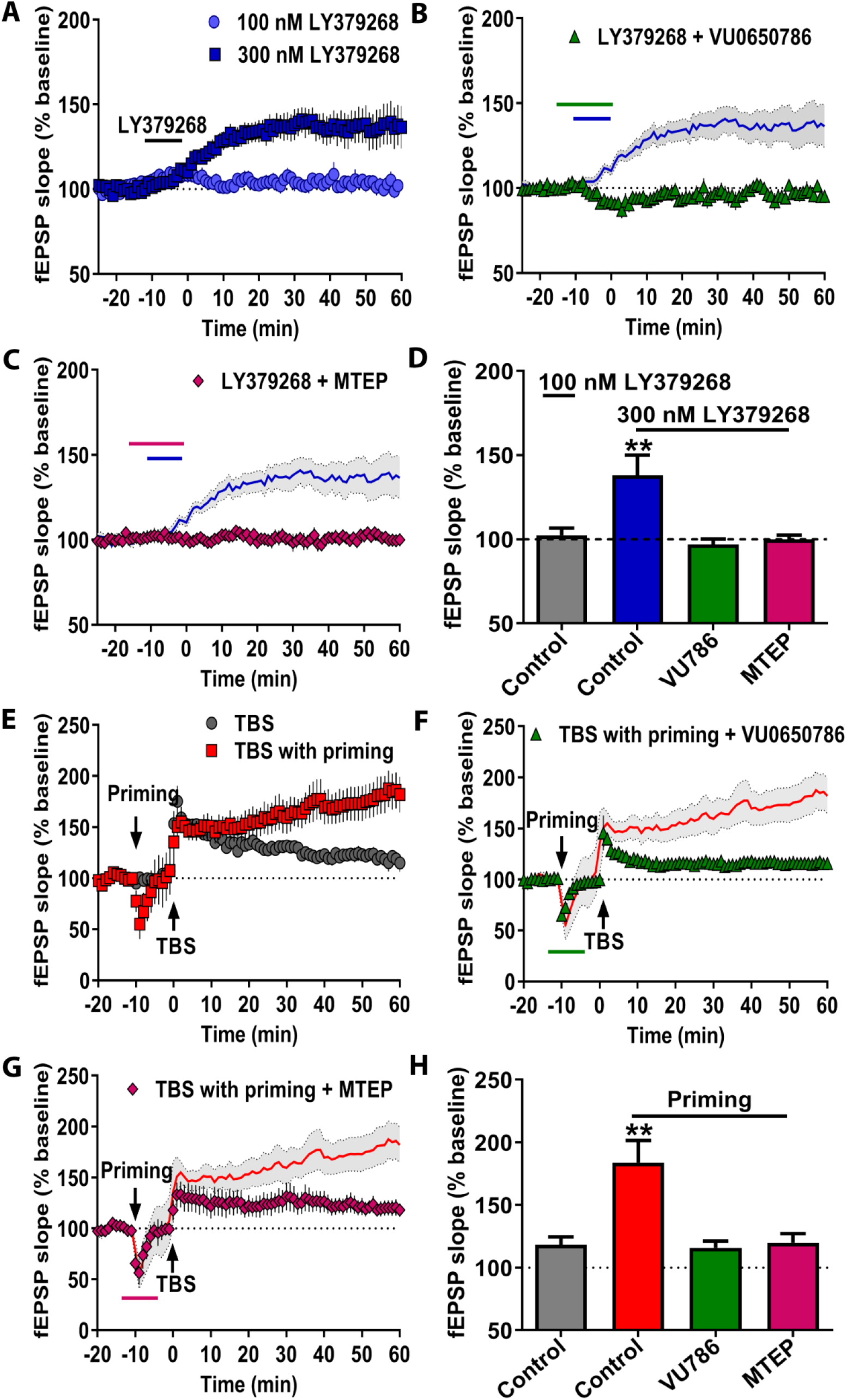
mGlu_3_ receptor signaling regulates multiple facets of LTP. (A) A high concentration of LY379268 (300 nM) and sustained 0.5 Hz stimulation enhanced fEPSP slope in acute hippocampal slices (dark blue squares, n=6 slices). A modest concentration of LY379268 (100 nM) did not affect fEPSP slope on its own (light blue circles, n=8). (B) The mGlu_3_ NAM VU0650786 (10 μM) blocked the fEPSP enhancement induced by LY379268 application (green triangles, n=5). Blue line displays LY379268 (300 nM) data from panel A. (C) The mGlu_5_ NAM MTEP (1 μM) blocked LY379268-induced LTP (pink diamonds, n=5). (D) Summary of averaged fEPSP slope of last 5 minutes of recordings from panels A-C (**p<0.01, compared to Control, F(3,20)=5.672, p<0.01 one-way ANOVA with Tukey’s post-hoc test). (E) Brief priming stimulation (2, 10-second, 10 Hz trains) followed by TBS (red squares, n=5) enhanced fEPSP slope compared to TBS alone (gray circles, n=5). (F) The mGlu_3_ NAM VU0650786 (10 μM) perfused before, during, and after priming stimulation blocked the enhanced LTP (green triangles, n=4). Red lines display TBS with priming data from panel E. (G) The mGlu_5_ NAM MTEP (1 μM) perfused throughout priming stimulation blocked the enhanced LTP (pink diamonds, n=4). (H) Summary of averaged fEPSP slope of last 5 minutes of recordings from E-G (**p<0.01, compared to Control, F(3,14)=9.136, p<0.01 one-way ANOVA with Tukey’s post-hoc test). Data are presented as mean ± SEM.

Interestingly, the effects of mGlu_3_ receptor activation outlined above are reminiscent of a previously reported form of mGlu_5_ receptor-dependent metaplasticity, referred to as LTP priming (23). Priming is a phenomenon in which modest activation of mGlu_5_ receptors with a brief stimulation of glutamatergic afferents exerts minimal acute effects on excitatory transmission but dramatically facilitates LTP induction by subsequent TBS afferent stimulation. We primed CA3-CA1 synapses with brief afferent stimulation (2 bursts of 10 stimuli at 10 Hz), and observed a robust increase in subsequent TBS-induced LTP relative to control slices (Figure 4E/4H). In light of the finding that mGlu_3_ activation potentiates TBS-LTP, we hypothesized that endogenous mGlu_3_ receptors might be involved in this process. Consistent with this hypothesis, the mGlu_3_ NAM VU0650786 blocked priming metaplasticity and returned TBS-LTP to control levels (Figure 4F/4H). In addition, consistent with previous studies (23), the mGlu_5_ NAM MTEP (Figure 4G/4H) also prevented the priming enhancement of TBS-LTP. Together, these results demonstrate that mGlu_3_ receptor activation induces this form of metaplasticity through coordinated signaling with mGlu_5_ receptors.

### mGlu_3_ receptor activation disinhibits CA1 pyramidal cells

mGlu_5_ receptor-dependent metaplasticity requires release of endocannabinoids, activation of CB1 endocannabinoid receptors, and the inhibition of GABAergic transmission onto pyramidal cells (23). We directly tested whether mGlu_3_ receptors induce similar CB1-dependent effects on inhibitory transmission by isolating evoked IPSCs (eIPSCs) on CA1 pyramidal cells. LY379268 (300 nM) acutely depressed eIPSCs in control recordings (Figure 5A/5D), which were blocked in the presence of the mGlu_3_ NAM VU0650786 (20 μM; Figure 5B/5D). The CB1 receptor antagonist AM251 (2 μM) also blocked the decrease in eIPSCs (Figure 5C/5D). Further, we returned to field potential recordings to test the hypothesis that activation of CB1 receptors is required for mGlu_3_ receptor-induced metaplasticity. AM251 (2 μM) co-application blocked LTP induced by high concentration LY379268 (Figure 5E/5F) and prevented effects on TBS-LTP (Figure 5G/5H). Together, these results suggest that mGlu_3_ receptors regulate hippocampal synaptic plasticity by recruiting endocannabinoid-mediated disinhibition downstream of postsynaptic mGlu receptors.

**Figure 5.**
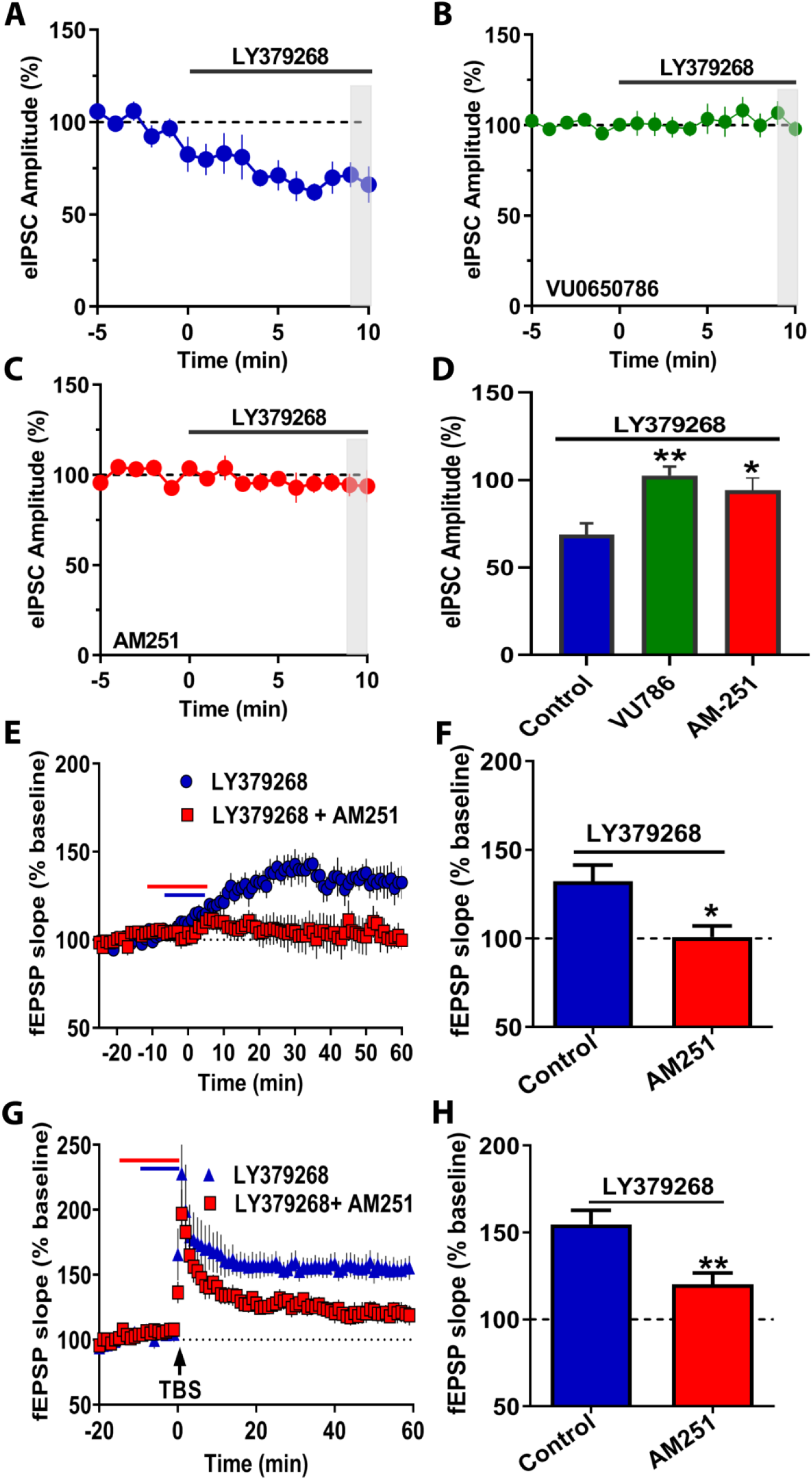
mGlu_3_ receptor activation decreases evoked inhibitory transmission onto hippocampal pyramidal neurons. (A) Electrically-evoked inhibitory postsynaptic currents (eIPSCs) were recorded from pyramidal cells in CA1. A high concentration of the mGlu_2/3_ agonist LY379268 (300 nM) reduced the amplitude of eIPSCs (blue circles, n=6 cells). (B) Co-application of the mGlu_3_ NAM, VU0650786 (20 μM) blocked the LY379268-induced decrease in eIPSCs (green squares, n=10). (C) The CB1 antagonist AM251 (2 μM) blocked the effects of LY379268 on eIPSC amplitude (red circles, n=6). (D) Summary of the last 2 min of the recordings from the time course experiments in panels A-C (*p<0.05, **p<0.01 compared to control, F(2,19)= 8.042, one-way ANOVA with Tukey’s post-hoc test). (E) In field potential configuration, AM251 (2 μM) blocked LTP (red squares, n=5 slices) induced by high concentration of LY379268 (300 nM) (blue circles, n=5). (F) Summary of averaged fEPSP slope of last 5 minutes of recordings from panel E (*p<0.05, compared to Control, t(8)=2.86, Student’s t-test). (G) AM251 (red squares, n=5) blocked the ability of LY379268 (100 nM) to enhance LTP following TBS (blue circles, n=5). (H) Summary of averaged fEPSPs slope of last 5 minutes of recordings from panel H. (**p<0.01, compared to Control, t(8)=3.617, Student’s t-test). Data are expressed as mean ± SEM.

### Neuronal mGlu_3_ receptors mediate hippocampal metaplasticity

Previous studies suggest that mGlu_5_-induced effects on hippocampal LTP are mediated by activation of mGlu_5_ expressed in hippocampal pyramidal cells (23). Based on this, the simplest hypothesis to explain our findings is that mGlu_3_ interacts with mGlu_5_ in hippocampal pyramidal cells to regulate LTP and trace fear conditioning.

However, the mGlu_3_ receptor is expressed in multiple neuronal populations and is heavily expressed in astrocytes, and it is possible that mGlu_3_ expressed in any of these cell populations could be required for enhancing LTP and trace fear conditioning. We therefore generated conditional *Grm3^Fl/Fl^* mice to genetically manipulate mGlu_3_ receptor expression (Figure 6A/6B and Supplementary Methods). We first validated the *Grm3^Fl/Fl^* construct by introducing Cre recombinase under the ubiquitously expressed cytomegalovirus (CMV) minimal promotor. We detected excision of *Grm3* exon 3 (Figure 6C), indicating Cre-mediated recombination readily occurs *in vivo*. We also performed RT-PCR, RNAscope, and western blots from hippocampal tissues and detected minimal transcript and protein in *Grm3*-CMV-KO mice (Figure 6D-6G).

**Figure 6.**
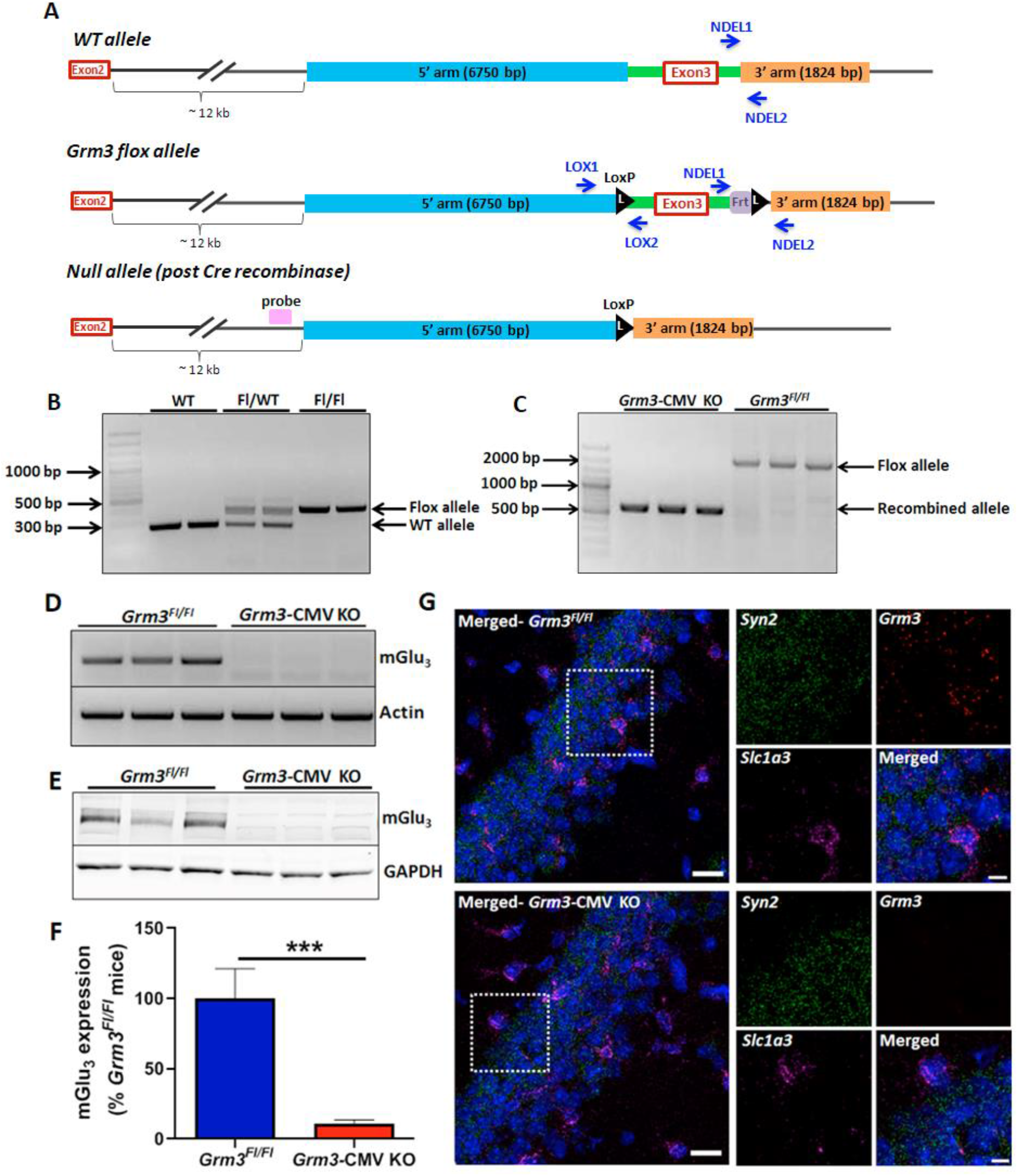
Generation and characterization of conditional Grm3^Fl/Fl^ mice. (A) Schematic of the procedure for generating the floxed *Grm3* clones. (Top) Wild-type *Grm3* locus surrounding exon 3. (Middle) Floxed *Grm3* locus where LoxP sites are inserted flanking exon 3 of *Grm3*. (Bottom) Cre-mediated recombination leading to loss of *Grm3* exon 3 and Frt site. (B) PCR from the DNA of mice homozygous for the WT allele (277 bp) and mice heterozygous and homozygous for the floxed allele (413 bp). CMV-Cre mediated recombination in *Grm3^Fl/Fl^* mice excises *Grm3* exon 3, leading to loss of WT allele (~1900 bp) and generation of a smaller recombination allele (671 bp). RT-PCR from the hippocampi of WT and *Grm3*-CMV KO mice. (E) Western blot depicting loss of mGlu_3_ protein from the hippocampus of *Grm3*-CMV KO mice (red bar) relative to *Grm3^Fl/Fl^* controls (blue bar). (F) Bar graph depicting quantification of Western blots. Data are presented as mean ± SEM, N=3-6 mice. (***p<0.001 compared to *Grm3^Fl/Fl^* mice, t(7)= 7.018, Student’s t-test). (G) Confocal 40X RNAscope *in situ* hybridization images showing loss of *Grm3* mRNA (red) from both neurons (*Syn2*; *Synapsin*, green) and astrocytes (*Slc1a3*; G*LAST*, magenta) in *Grm3*-CMV KO mice. Scale bar = 20 μm for the merged left panel image, and 10 μm for the 3X images.

To test the hypothesis that that postsynaptic mGlu_3_ receptors on pyramidal cells are required for enhancing hippocampal synaptic plasticity, we crossed *Grm3Fl*/*Fl* with CaMKII Cre mice, a selective marker for glutamatergic neurons in the cortex and hippocampus (23, 42) and performed experiments at an age that displays preferential receptor ablation from the CA1 area of hippocampus (Figure 7A). We assessed cell type-specificity using RNAscope and observed *Grm3* ablation from cells co-expressing transcript for vGluT1 (*Slc17a7*) but not GLAST (*Slc1a3*) in hippocampus from *Grm3*-CaMKII KO mouse (Figure 7B). CaMKII Cre mice display recombination in the striatum during late adulthood (43); however, we observed no changes in *Grm3* in the nucleus accumbens of 6-8-week-old *Grm3-*CaMKII KO mice (Figure S3). We performed similar experiments to validate *Grm5* deletion from *Grm5-*CaMKII KO mice (Figure S4) (see also (23)).

**Figure 7.**
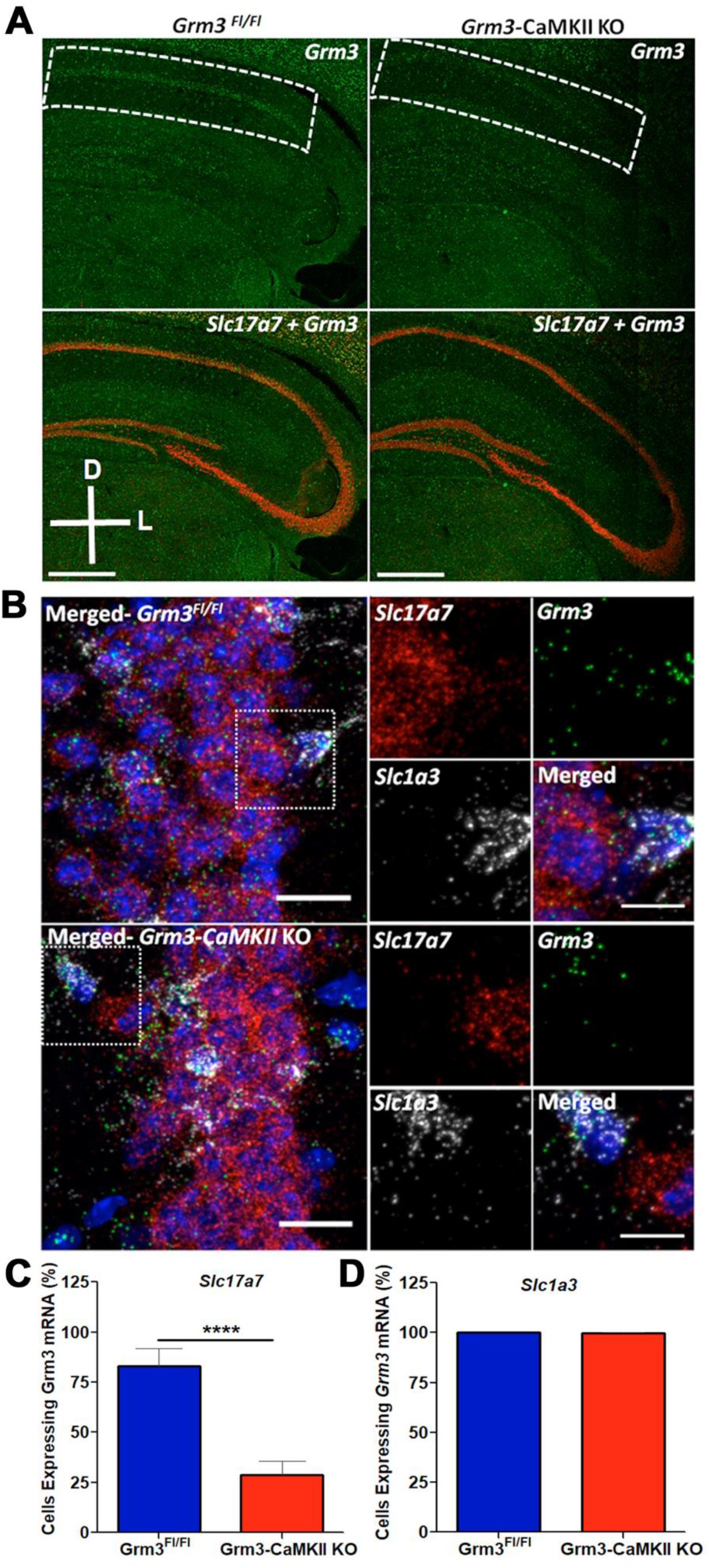
Grm3-CamKII KO mice display selective ablation of mGlu_3_ receptor transcript in pyramidal cells of the hippocampus. (A) Characterization of *Grm3-*CamKII KO mice. Merged confocal 20X tile scan image of a coronal section highlighting decreased *Grm3* mRNA expression in the CA1 of 7-8-week-old *Grm3*-CaMKII KO mice. Scale bars denote 500 μm. (B) Representative confocal 40X RNAscope *in situ* hybridization images showing loss of *Grm3* mRNA (green) from pyramidal neurons (*Slc17a7*; vGluT1, red) of CA1, while *Grm3* mRNA expression is intact in astrocytes (*Slc1a3*; GLAST, gray). Scale bars denote 20 μm for the left panel image and 10 μm for the right panels (per genotype: *Grm3^Fl/Fl^*=3; *Grm3-* CaMKII KO=4). (C) The percentage of Slc17a7-positive cells (vGluT1) with *Grm3* mRNA was decreased in *Grm3-*CaMKII KO mice (red bar, n/N = 16/4 slices/mice) as compared to WT controls (blue, barn/N = 15/3) (****p<0.0001 compared to *Grm3^Fl/Fl^* mice, t(29) = 19.40, Student’s t-test). (D) The percentage of Slc1a3-positive cells (GLAST) with *Grm3* mRNA was not different between *Grm3-*CaMKII KO mice (red bar, n/N = 14/4) and controls (blue bar, n/N = 9/3) (t(21) = 0.7951, Student’s t-test, p=0.4355).

With these new transgenic tools, we proceeded to test the hypothesis that neuronal mGlu_3_ receptors mediate hippocampal metaplasticity. Consistent with our hypothesis, LY379268 (100 nM) potentiated TBS-LTP in slices from Cre- (littermate controls) but not Cre+ (*Grm3*-CaMKII KO) mice (Figure 8A/8B), suggesting mGlu_3_ receptors expressed in the pyramidal neurons are necessary for LTP enhancement. As hippocampal metaplasticity is important for associative learning (23), we reasoned that neuronal mGlu_3_ receptors would be essential for cognitive enhancement following mGlu_3_ receptor activation. To test this hypothesis, we performed trace fear conditioning experiment using *Grm3*-CaMKII KO mice and littermate controls. We observed increased freezing in female *Grm3*-CaMKII KO mice relative to controls (data not shown), therefore, we only used male *Grm3*-CaMKII KO mice to examine potentiation by LY379268. Systemic LY379268 administration potentiated trace fear conditioning in littermate control mice, however no difference in freezing was observed between Vehicle- and LY379268-treated *Grm3*-CaMKII KO male mice (Figure 8C). We obtained similar physiological and behavioral results using *Grm5*-CaMKII KO mice (Figure 8D-8F), indicating that concerted interactions between neuronal mGlu_3_ and mGlu_5_ receptors mediate hippocampal metaplasticity and pro-cognitive effects. In sum, the data obtained from transgenic mice corroborate our pharmacological findings and provide new evidence that mGlu_3_ receptor activation enhances cognition through direct actions on hippocampal pyramidal cells.

**Figure 8.**
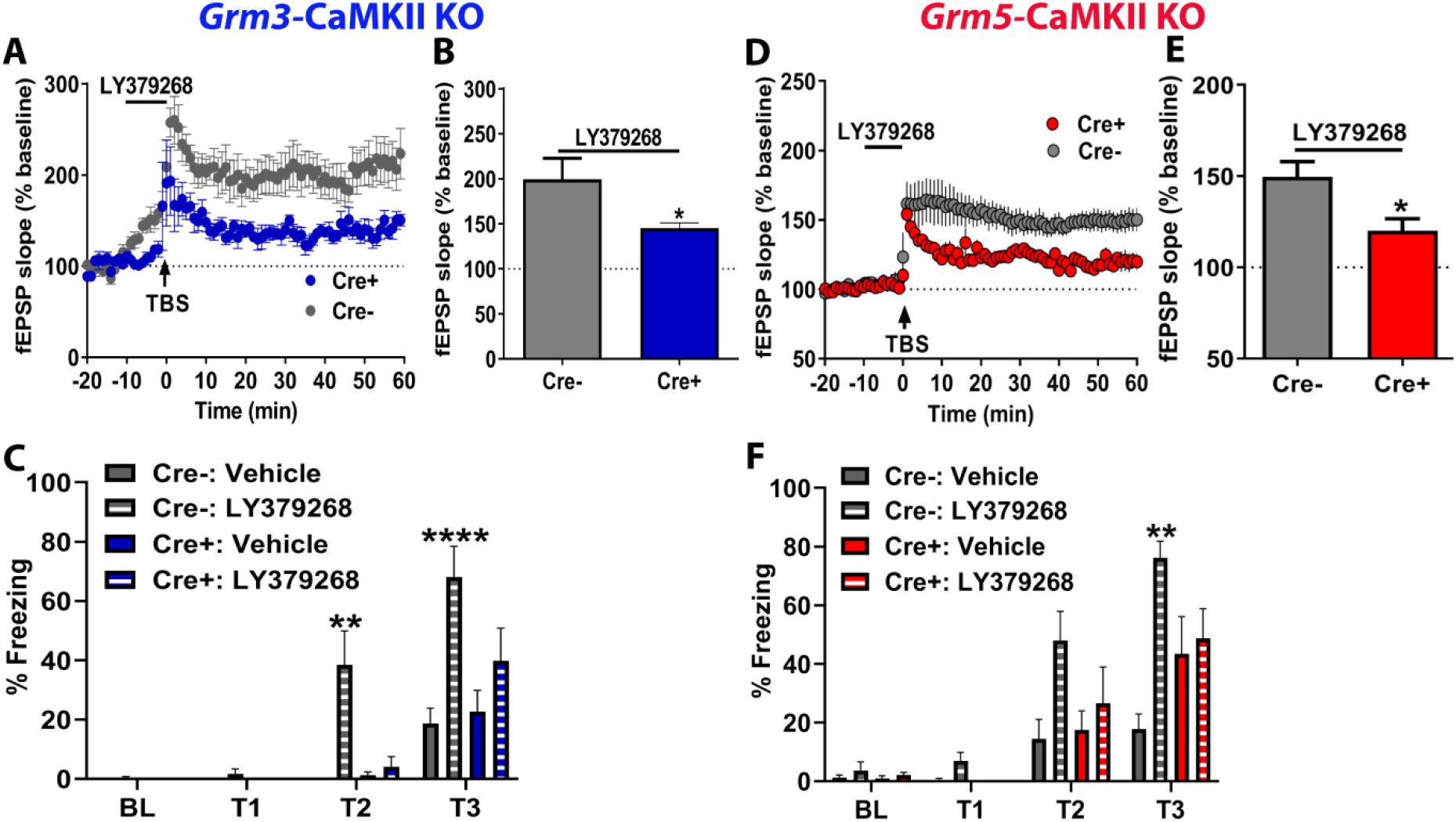
Neuronal mGlu receptors mediate cognitive enhancement following mGlu_3_ receptor activation. (A) LY379268 (100 nM) failed to enhance LTP after TBS in slices from Cre+ knockout mice (blue circles, n=6) relative to Cre-littermate controls (grey circles, n=5). (B) Summary of averaged fEPSPs of last 5 minutes of recordings from panel C (*p<0.05 compared to Cre-, t(9)= 2.506, Student’s t-test). (C) Systemic treatment with LY379268 (3 mg/kg) enhances the expression of trace fear conditioning in Cre-littermate controls (hatched grey bars, n=6 mice) but not in Cre+ knockout mice (hatched blue bars, n=6 mice) as compared to treatment with vehicle (gray bars, Cre-, N=3 mice; blue bars, Cre+, N=3 mice) (**p<0.01, ****p<0.001 compared to Cre-Vehicle, F(3,56)=23.41, two-way repeated measures ANOVA with Tukey’s post-hoc test). (D) LY379268 (100 nM) potentiates LTP after TBS stimulation in Cre-controls (grey circles, n=5) but not Cre+ slices (red circles, n=5). (E) Summary of averaged fEPSPs of last 5 minutes of recordings (*p<0.05 compared to Cre-, t(8)= 2.7, Student’s t-test). (F) Loss of enhanced freezing expression following LY379268 treatment (hatched bars) in Cre+ mice (red hatched bars, 7 mice) as compared to Cre-littermate controls (grey hatched bars, 8 mice) (**p<0.01 compared to Cre-Vehicle, F(3,72)=2.788 by two-way repeated measures ANOVA with Tukey’s post-hoc test, N=4-9 mice). Data are expressed as mean ± SEM.

## Discussion

Genetic studies have revealed associations between *GRM3* variations and cognitive function in patients with schizophrenia, substance abuse disorders, major depression, and autism (2, 5, 44, 45). Despite these links, the hypothesis that mGlu_3_ receptors promote cognition has not been evaluated in mechanistic preclinical studies. Here, we demonstrated that activating mGlu_3_ receptors enhances associative learning in mice and in a model of schizophrenia-like cognitive dysfunction. We then found that hippocampal mGlu_3_ receptor activation rescues plasticity deficits by recruiting mGlu_5_ receptor-dependent, endocannabinoid-mediated disinhibition. Finally, we generated a new line of transgenic mice and found that mGlu_3_ receptors that are co-expressed with mGlu_5_ receptors in hippocampal projection neurons are required for the cognitive enhancing properties of mGlu_3_ receptor activation.

Multiple mGlu receptors modulate hippocampal physiology (3), but assigning specific roles to individual receptor subtypes has been hampered by low availability of tools to differentially modulate their functions. We recently discovered molecular probes that comprise the first selective mGlu_3_ receptor ligands (20, 46). Using these tools and global knockout mice, we found that mGlu_3_ receptors regulate synaptic plasticity in the prefrontal cortex through pathways that are also regulated by mGlu_5_ receptors (30, 31). Here, we observed a similar functional interaction between mGlu_3_ and mGlu_5_ receptors in area CA1 of the hippocampus. This is an especially interesting finding in light of extensive previous studies suggesting that mGlu_5_ receptors bidirectionally modulate hippocampal synaptic plasticity (23, 32, 36) and improve multiple aspects of hippocampal-dependent cognitive function (36, 47-49). Interestingly, mGlu_5_ receptors promote LTP and LTD through distinct mechanisms: LTP potentiation requires calcium-dependent endocannabinoid-mediated disinhibition (23, 37), whereas phosphoinositide-3-kinase (PI3K), Akt, and mTOR signaling initiate LTD (50). The present finding that mGlu_3_ activation selectively enhances LTP but not LTD suggests that that hippocampal mGlu_3_ receptors may bias mGlu_5_ signaling away from the PI3K cascade that induces LTD. These findings contrast with observations in prefrontal cortex pyramidal cells, where mGlu_3_ receptors direct synaptic plasticity through mGlu_5_-PI3K-Akt signaling (30). This distinction between hippocampal and neocortical plasticity begs the question: what are the molecular determinants that guide the rules of mGlu receptor-dependent synaptic plasticity? Homer proteins comprise an extremely likely candidate family, as their interactions with mGlu_5_ are required for both hippocampal mGlu_5_-LTD (50) and PFC mGlu_3_-LTD (30), and they can bias mGlu_5_ receptors towards specific signaling pathways (51, 52). Interactions between mGlu receptors and NMDA receptors and other synaptic proteins are likely to shape the metaplastic landscape as well and merit further investigation.

mGlu_3_ receptors recruit a distinct component of mGlu_5_ receptor signaling to modulate hippocampal function. This finding has exciting ramifications given the abundance of evidence indicating mGlu_5_ positive allosteric modulators (PAMs) enhance cognition in preclinical models of pathophysiological changes thought to be important for symptoms of schizophrenia (3). mGlu_5_ PAMs have not reached clinical trials and multiple companies have halted efforts to advance mGlu_5_ PAMs to clinical development. One contributing factor was concern that stemmed from excitotoxicity induced by specific, structural classes of related mGlu_5_ modulators that have allosteric agonist activity and potentiate NMDA receptor currents. Recent efforts have generated new scaffolds of biased mGlu_5_ PAMs that do not have allosteric agonist activity or enhance NMDA receptor currents but retain the ability to modulate discrete aspects of mGlu_5_ actions in the CNS (21, 30, 37, 47, 53). Based on the effects of mGlu_3_ activation outlined in the current studies, it is possible that selective activators of mGlu_3_ could provide an alternative approach to mGlu_5_ PAMs for enhancing multiple domains of cognitive function. Thus, it will be important to focus major efforts on discovery of highly selective mGlu_3_ agonists and PAMs that can be used to fully evaluate their potential in alleviating cognitive disturbances in preclinical models of schizophrenia. However, like mGlu_5_ receptors (37, 47, 54), previous studies suggest that mGlu_3_ receptors may directly regulate NMDA receptor currents in young animals (55), but mGlu receptors display high constitutive activity early in development (31, 56) and it remains unclear whether a direct mGlu_3_-NMDA receptor interaction persists into adulthood. Regardless, astrocytic mGlu_3_ receptors exert neuroprotective effects (57, 58), providing an additional safeguard against potential excitotoxicity and further rationale for developing mGlu_3_ PAMs as safer alternatives to other investigational cognitive enhancers.

Collectively, these pharmacological and genetic studies provide convergent and concrete evidence for the cellular location of mGlu_3_ receptors responsible for regulating hippocampal LTP and trace conditioning. Previous studies indicate mGlu_3_ receptors are heavily expressed in astrocytes and in pre- and post-synaptic structures of neurons (59, 60). The current results showed no alterations in paired-pulse ratio during LTP and demonstrated that postsynaptic signaling is required for mGlu_3_-mediated disinhibition. Furthermore, the effects of mGlu_3_ activation are absent in mice in which mGlu_3_ or mGlu_5_ receptors have been selectively deleted from pyramidal cells. Together, these data demonstrate that the ability of mGlu_3_ activation to direct hippocampal plasticity and related behavior is mediated by receptor populations interacting postsynaptically on CA1 pyramidal cells. Nonetheless, astrocytic mGlu_3_ receptors are likely to regulate neurotransmission and modulate other aspects of disease-relevant physiology. Future research utilizing cell type-specific manipulations of mGlu_3_ receptor function will improve our understanding of how discrete receptor populations regulate neurophysiology and guide behavior.

Hippocampal lesions generate profound impairments of trace conditioning in patients with amnesia and in rodent models (61–63). Here, conversely, we observed enhanced trace fear conditioning following hippocampal modulation by mGlu_3_ receptor activation. Furthermore, these findings provide evidence for a causative association, as the pro-cognitive effect required mGlu receptors expressed in hippocampal pyramidal cells. We performed experiments with young adult *Grm3*-CaMKII KO and *Grm5*-CaMKII KO mice, a timepoint that mitigate extrahippocampal receptor ablation (23, 42). Despite this design, the possibility that decreased expression of extrahippocampal receptors contributed to the behavioral changes in knockout mice should cannot be excluded, as mGlu_3_ receptors within the prefrontal cortex regulate synaptic plasticity (26, 30) and the neocortex is involved in components of trace conditioning (64). Nonetheless, our findings demonstrate that mGlu_3_ receptors recruit hippocampal circuits and downstream mechanisms that have been intimately linked with trace fear conditioning (23, 65, 66). For one, genetic reduction of hippocampal GABAA receptor expression enhances trace fear conditioning (66), suggesting that CA1 disinhibition is sufficient to enhance associative learning. Furthermore, we observed that mGlu_3_ activation reversed cognitive deficits in mice treated with scPCP. Previous studies have revealed that scPCP treatment enhances inhibitory tone onto CA1 pyramidal cells and thereby raises the threshold for LTP induction (28). We therefore posit that mGlu_3_ agonism facilitated LTP induction in scPCP hippocampal slices by restoring inhibition, thereby reversing trace conditioning deficits in the model of schizophrenia-like forebrain disruption.

Collectively, these combined electrophysiological and behavioral studies highlight that mGlu_3_ receptors regulate hippocampal plasticity and promote associative learning. Further translational studies will be needed to fully scrutinize this hypothesis, but the current results suggest that mGlu_3_ receptors are well-suited for ameliorating cognitive symptoms in schizophrenia.

## Supporting information

Key Resource Table

## Acknowledgements

The authors would like to thank Dr. Margarita Behrens for providing the initial breeders for the *Grm5^Fl/Fl^* mice used in these studies and Dr. Xiaoyan Zhan and Dr. Hyekyung Plumley for assistance establishing the *Grm3^Fl/Fl^* mice. Behavioral experiments were performed through the Murine Neurobehavior Core lab at the Vanderbilt University Medical Center. Confocal imaging was performed in part through the Vanderbilt Cell Imaging Shared Resource (supported by NIH grants CA68485, DK20593, DK58404, DK59637 and EY08126). This work was supported by NIH grants F32MH111124 (B.J.S), K01MH112983 (R.G.G.), K99AA027806 (M.E.J.), R01MH062646 (P.J.C.), and R37NS031373 (P.J.C.) and rettsyndrome.org grant #3503 (C.M.N).

## Author contributions

Conceptualization, S.D., B.J.S., F.N., M.E.J., P.J.C.; Methodology, S.D., B.J.S., Z.X., R.G.G, M.E.J.; Validation, S.D.; Investigation, S.D., B.J.S., Z.X., W.Q.; Resources, C.W.L., P.J.C.; Writing – Original Draft, S.D., M.E.J.; Writing – Review and Editing, all authors; Supervision, C.M.N., M.E.J., P.J.C.

