## Supplementary material for "Activating mGlu_3_ metabotropic glutamate receptors rescues schizophrenia-like cognitive deficits through metaplastic adaptations within the hippocampus": Key Resource Table

**Key Resources Table**

| **Reagent or Resource** | **Source** | **Identifier** |
| --- | --- | --- |
| **Antibodies** | | |
| rabbit anti-mGlu_3_ antibody (1:500) | Alomone | AGC-012 |
| mouse anti- GAPDH antibody (1:5000) | ThermoFisher | MA5-15738 |
| goat anti-rabbit 800 (1:5000) | LiCor | 925-32211 |
| goat anti-mouse 680 (1:5000) | LiCor | 926-68070 |
| **RNAscope *in situ* Hybridization Probes** | | |
| *Grm3* (target region 875-1676) | ACDBio | Mm-Grm3-O (NM_181850.2) |
| *Grm5* (target region 2409-3336) | ACDBio | Mm-Grm5-O1 (NM_001081414.2) |
| *Slc17a7* (target region 464-1415) | ACDBio | Mm-Slc17a7-C3 (NM_182993.2) |
| *Syn2* (target region 585-1482) | ACDBio | Mm-Syn2-C2 (NM_013681.3) |
| *Slc1a3* (target region 1122-2237) | ACDBio | Mm-Slc1a3-C3 (NM_148938.3) |
| *Slc32a1* (target region 894-2037) | ACDBio | Mm-Slc32a1-C2 (NM_009508.2) |
| **Chemicals** | | |
| LY379268 (3 mg/kg; 100-300 nM) | Tocris | 5064 |
| D-AP5 (50 µM) | Tocris | 0106 |
| DHPG (25-50 µM) | Tocris | 0805 |
| CNQX (20 µM) | Tocris | 1045 |
| MTEP (1 µM) | Tocris | 2921 |
| AM251 (2 µM) | Tocris | 1117 |
| Phencyclidine (10 mg/kg) | Sigma | P3029 |
| VU0469650 (10 µM) | CW Lindsley (Lovell et al., 2013) | N/A |
| VU0650786 (30 mg/kg; 20 µM) | CW Lindsley (Engers et al., 2017) | N/A |
| VU6001966 (10 mg/kg; 10 µM) | CW Lindsley (Bollinger et al., 2017) | N/A |
| **Experimental Models** | | |
| C57BL/6J | The Jackson Laboratory | 000664 |
| *Grm5^Fl/Fl^* | The Jackson Laboratory | 028626 |
| *Grm3^Fl/Fl^* | See Methods | N/A |
| CaMKII Cre | The Jackson Laboratory | 005359 |
| CMV Cre | The Jackson Laboratory | 006054 |
