## Supplemental info for "Activating mGlu_3_ metabotropic glutamate receptors rescues schizophrenia-like cognitive deficits through metaplastic adaptations within the hippocampus"

**Supplementary Methods**

**RNAscope *in situ* hybridization (ISH) and confocal imaging.** Brains were rapidly dissected and flash-frozen on dry ice within 5 minutes of harvest. Frozen tissue was embedded in cryo-embedding medium and stored at -80°C until sectioning for ISH. Brain slices (16 µm) containing the desired brain areas were obtained utilizing a Leica CM 3050S cryostat at -20°C, mounted on frosted microscope slides, and stored at -80°C until processing. The following steps were performed at room temperature unless otherwise noted. Briefly, slides were fixed in 4% paraformaldehyde for 20 minutes at 4°C, immediately followed by dehydration with 50% ethanol (5 minutes), 70% ethanol (5 minutes), and twice with 100% ethanol (5 minutes). After dehydration, slides were dried for 5 minutes on absorbent paper. A hydrophobic barrier was drawn around the sections using a hydrophobic pen and allowed to dry for 5 minutes. Sections were then incubated with Protease IV solution for 30 minutes and washed twice with PBS. Following wash, brain slices were incubated for 2 hours at 40°C with probes for cellular subtype markers and with probes we designed to recognize the gene sequence targeted for excision in the *Grm3^Fl/Fl^* and *Grm5^Fl/Fl^* mice (Key Resources Table). All probes were purchased from ACDbio. Slices were washed twice with wash buffer before each amplification step. We incubated slides at 40°C with Amplification Reagents- Amp 1-FL (30 minutes), Amp 2-FL (15 minutes), AMmp 3-FL (30 minutes), and Amp 4 Alt A-Fl(15 minutes). Two washes for two minutes with 1X wash buffer were performed after each Amplification reagent application and finally, excess liquid was removed, and DAPI solution was applied to sections for 30 seconds. Slides were then covered with 1-2 drops of fluorescent mounting medium and stored at 4°C until imaged on a confocal microscope (Zeiss LSM 710). All images were collected with xyz acquisition mode where the total thickness of sample ranged between 5-7 μm. All the optical sections were projected as a single image using Zen 2.6 software and fluorescence signals were quantified using Imaris analysis and Imagej software. Cells with more than 2 puncta of *Grm3* mRNA were counted positive for *Grm3* expression and representative images were processed in Adobe Photoshop CS5.1.

**Generation of floxed *Grm3* mice.** The floxed *Grm3* mouse line was generated using FLP embryonic stem (ES) cell-mediated gene targeting on the C57BL/6J genetic background (Ingenious Targeting Laboratory). In these mice, exon 3 of *Grm3* is flanked by LoxP sites (Figure 6). To generate these mice, *Grm3* target vector was introduced into FLP ES lines carrying a FLP transgene, and the ES cell clones were screened to identify targeted clones in which FLP recombinase had deleted the neomycin selection cassette. The ES cells containing the selected clones of target *Grm3* allele were microinjected into Balb/c blastocysts, resulting in chimeras that were mated to C57BL/6J wild-type mice to generate “germline Neo deleted mice”. Tail DNA was screened for deletion of Neo cassette using primers specific for NDEL1: 5’-TACATGAAACTGAAAGGCCCCTAGG-3’ and NDEL2: 5’-TCCTATTCTGTCCCTAATTAGGCTCTGGG-3’ at the following amplification conditions: initial denaturation 95°C for 2 minutes, (denaturation 95°C for 30 seconds, annealing 60°C for 30 seconds, extension 72°C for 30 seconds) repeated 30 times.. Gel electrophoresis of the PCR product yielded a 277 bp amplicon for WT allele and a second amplicon of size 413 bp representing the floxed *Grm3* allele (Figure 6). All PCR products were sequence-verified for the presence of distal LoxP sites using Lox1: 5’-AAAAACAAAGCAAAGAAAAAGTTACTG-3’ and Lox2: 5’-CGTGTAGAGGTCGGCTTTTG-3’ specific primers, and following amplification conditions: initial denaturation 95°C for 2 minutes, (denaturation 94°C for 30 seconds, annealing 55°C for 30 seconds, extension 72°C for 60 seconds) repeated 30 times, hold at 4°C until further use. Mice positive for the *Grm3* floxed allele were mated to each other to generate homozygous mice (*Grm3^Fl/Fl^*).

**Supplemental Figure S1**

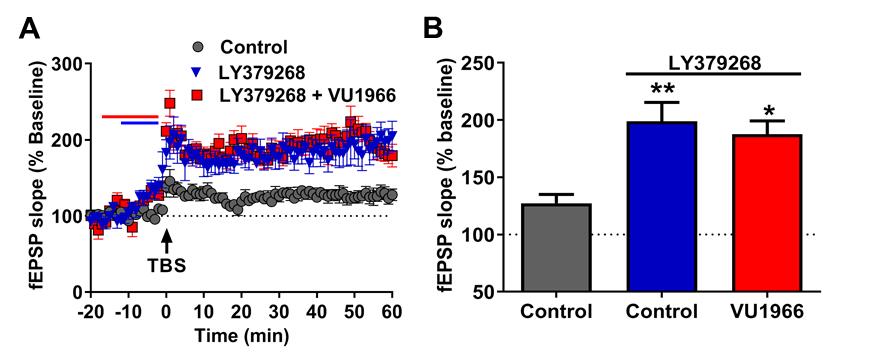

**Figure S1. mGlu2 NAM VU1966 does not block LY379268- induced long-term potentiation (LTD).at SC-CA1 synapse.**

(A) LY379268 application enhanced LTP in response to TBS (blue triangles, n=8 slices) as compare to control slices (gray circles, n=6 slices). Co-application of mGlu_2_ NAM VU1966 (10 µM) blocked the enhanced LTP induced by LY379268 (red squares, n=11). (B) Summary of averaged fEPSP slope of last 5 minutes of recordings from panel A (*p<0.05, ** p<0.01 compared to Vehicle, F_(2,22)_=6.976, one-way ANOVA with Tukey’s post-hoc test).

**Supplemental Figure S2**

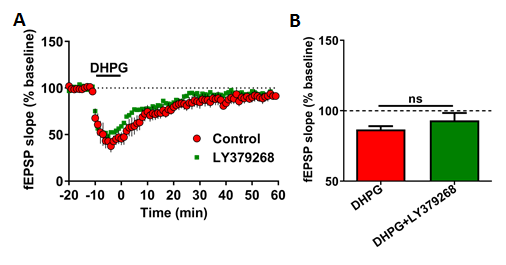

**Figure S2. LY379268 application does not alter threshold DHPG long-term depression (LTD).**

(A) Application of a threshold concentration of the mGlu_1/5_ agonist DHPG (25µM) for 10 minutes induced modest LTD of field excitatory postsynaptic potential (fEPSP) slope (n=4 slices). Co-application of the mGlu_2/3_ agonist LY379268 (100nM) for 10 minutes did not alter DHPG-induced LTD (n=6). (B) Summary of last 5 minutes of experiments (t_(8)_=0.9456, p=0.3720, Student’s t-test). Data are presented as mean ± SEM.

**Supplemental Figure S3**

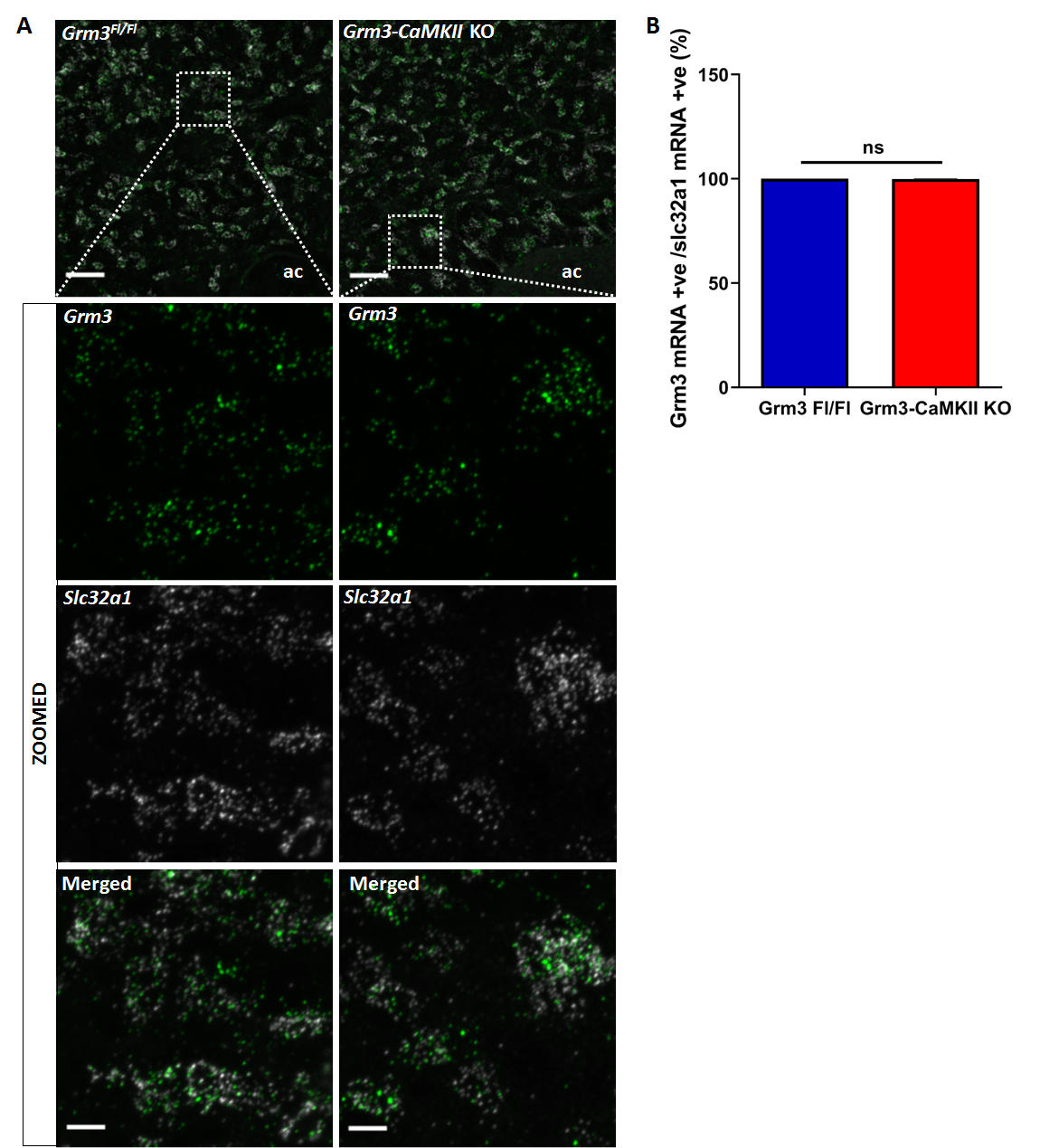

**Figure S3. Intact *Grm3* expression in the nucleus accumbens of *Grm3*-CamKII KO mice.**

**(A)** Representative confocal 20X RNAscope *in situ* hybridization images displaying no change in *Grm3* expression (green) in GABAergic neurons (*Slc32a1*; Vesicular GABA Transporter (VGAT); gray) within the nucleus accumbens of 6-8-week-old *Grm3*-CaMKII KO mice (mice per genotype: *Grm3^Fl/Fl^*=3; *Grm3-*CaMKII KO=4). Scale bar = 50 µm for the top panel image and 10 µm for the 5X zoomed images. **(B)** The percentage of *Slc32a1*-positive cells with *Grm3* mRNA did not change in *Grm3*-CaMKII KO mice (n/N=6/2 slices/mice) as compared to controls (n/N=5/2) (t_(9)_ = 1.414; Student’s t-test, p=0.1910).

**Supplemental Figure S4**

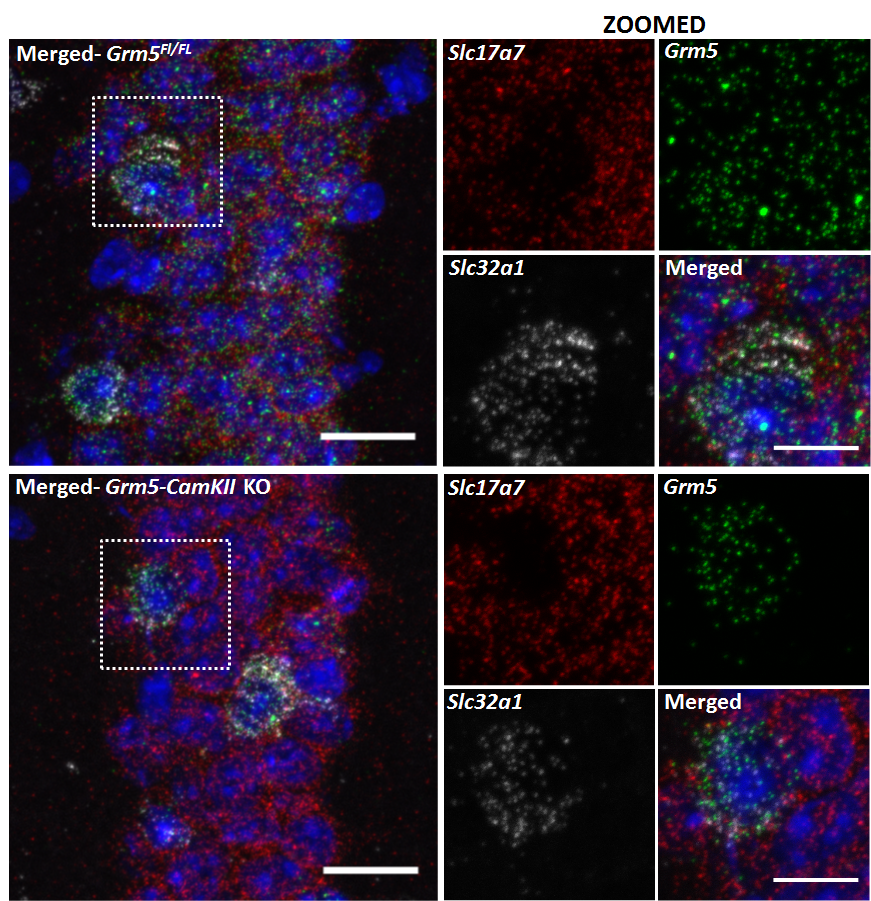

**Figure S4. Selective deletion of *Grm5* from glutamatergic neurons in *Grm5-*CaMKII KO mice**.

Representative confocal 40X RNAscope *in situ* hybridization images of displaying loss of *Grm5* (green) expression in glutamatergic neurons (*Slc17a7*; Vesicular Glutamate Transporter, VGluT, red) within the CA1 from 6-8-week old *Grm5*-CaMKII KO mice. *Grm5* expression in GABAergic neurons (*Slc32a1*; VGAT; gray) from *Grm5-*CaMKII KO mice is similar to littermate controls (mice per genotype: *Grm5^Fl/Fl^*=3; *Grm5-*CaMKII KO=3). Scale bar = 20µm for the merged left image and 10µm for the 3X zoomed images.
